# Increased mannosylation of extracellular vesicles in Long COVID plasma provides a potential therapeutic target for *Galanthus nivalis* agglutinin (GNA) affinity resin

**DOI:** 10.1101/2025.11.21.689519

**Authors:** Miguel A. Pesqueira Sanchez, Rosalia de Necochea Campion, Thomas Dalhuisen, Emily A. Fehrman, Pahul S. Chhabra, J. Daniel Kelly, Jeffrey N. Martin, Steven G. Deeks, Timothy J. Henrich, Michael J. Peluso, Steven P. LaRosa

**Affiliations:** Aethlon Medical, Inc., San Diego, CA, USA; University of California, San Francisco, San Francisco, CA, USA

## Abstract

There is no proven therapy for Long COVID, a post-acute illness characterized by a myriad of diverse symptoms including fatigue, dyspnea, and brain fog following SARS-CoV-2 infection. Extracellular vesicles (EVs) have been implicated in Long COVID pathogenesis by promoting viral and inflammatory signaling with their molecular cargo. In this study, we investigated whether EV abundance and glycome characteristics are altered in plasma from people with Long COVID and whether they can be targeted for removal using a glycan-binding affinity resin. Large (100−500 nm) and small (40−200 nm) EVs were isolated from plasma of participants in the post-acute phase of COVID-19 and analyzed by nanoparticle flow cytometry to measure concentration and glycan characteristics. Plasma of those with Long COVID contained elevated levels of both large and small EVs, and mannose-positive large EVs were significantly increased in comparison to recovered controls (p < 0.05). EV capture assays using *Galanthus nivalis* agglutinin (GNA) affinity resin demonstrated small EV removal positively correlated with mannose-positive EV abundance (r = 0.341, p < 0.05). NanoString analyses identified seven EV-associated miRNAs significantly depleted by GNA affinity resin treatment of plasma. PROGENy pathway inference of validated miRNA-mRNA interactions suggests these reductions may lead to a downregulation of JAK-STAT signaling and upregulation of Estrogen, VEGF, and PI3K pathways, resulting in a favorable rebalancing of immune and tissue-repair networks. These findings reveal specific glycome EV-miRNA cargo signatures in Long COVID and the potential clinical benefits of a lectin capture therapeutic strategy to remove these pathogenic vesicles and their inflammatory cargo.

## Introduction

Following the acute phase of SARS-CoV-2 infection, a subset of people experience Long COVID [1], a post-acute condition characterized by ongoing symptoms that may include fatigue, post-exertional malaise, dyspnea, chest pain, and troubling neuro-psychiatric problems including “brain fog” [2, 3]. Despite its estimated prevalence— 6-7% of the U.S. population [4], potentially hundreds of millions of people globally and its large burden (∼1 trillion dollars per year [5]), management remains focused on symptom relief and no curative treatments are available.

Multiple pathophysiologic mechanisms have been proposed to contribute to the development of Long COVID. These include persistence of SARS-CoV-2 RNA and protein, reactivation of latent herpesviruses (e.g., Epstein-Barr Virus), immune dysregulation and autoimmunity, thrombosis, dysbiosis, and mitochondrial and tissue dysfunction [6, 7]. Extracellular vesicles (EVs), small nanoparticles with a lipid bilayer approximately 30−1000 nm in diameter, released from all cell types and involved in cell-to-cell communication, have been implicated in the pathogenesis of Long COVID [8]. EVs have been found to contain viral components including the SARS-CoV-2 spike protein [9, 10].

EV surface glycans, including high-mannose N-glycans, regulate vesicle biogenesis, biodistribution, cellular uptake, and immune interactions, making EV glycosylation a central functional feature and potential target for capture or diagnostics [11]. In acute COVID-19, there is evidence supporting a functional glycome mechanism in disease pathology, with glycan changes observed in immune effector proteins in both plasma and lung tissues [12]. Notably, the Aethlon Hemopurifier®, which targets specific glycan structures, was shown to remove circulating SARS-CoV-2 virus, extracellular vesicles, and pathogenic miRNA cargo from critically ill COVID-19 patients [13]. The Hemopurifier® is a plasmapheresis cartridge filled with *Galanthus nivalis* agglutinin (GNA) affinity resin, which binds mannosylated glycoproteins found on the surface of enveloped viruses and some EVs [14]. Persistent glycome remodeling of serum proteins in acute COVID-19 has been associated with disease severity and duration in multiple independent cohorts [15], raising questions about whether similar alterations persist in Long COVID plasma-derived EVs.

In this study, we measured the abundance and mannosylation characteristics of plasma-derived EVs from participants with Long COVID and compared them to EVs from individuals who had fully recovered after an episode of COVID-19. Our objective was to determine whether increased EV production or mannose glycosylation are traits of this illness. We describe the association between Long COVID EV mannosylation patterns and capture dynamics on the GNA affinity resin. Furthermore, we examine the miRNA cargo of GNA-captured Long COVID EVs to gain insight into their potential contributions to Long COVID pathogenesis. Pathway inference analysis allows us to identify changes in core signaling networks modulated by miRNA-mRNA interactions implicated in immune dysregulation in Long COVID. This novel research broadens our understanding of how individuals afflicted with Long COVID may benefit from strategies such as the Aethlon Hemopurifier® to remove pathogenic EVs and miRNA cargo.

## Materials and Methods

### Participant selection and evaluation

All samples were from individuals enrolled in the San Francisco-based Long-term Impact of Infection with Novel Coronavirus (LIINC) cohort (NCT04362150). Details of the cohort design have been previously published [16]. Briefly, adults with a history of test-confirmed SARS-CoV-2 infection are assessed at baseline and at approximately 3-month intervals thereafter, during which they undergo interviewer-administered assessments of COVID-attributed symptoms, quality of life, and medical comorbidities, as well as biospecimen collection. Following clinician review, participants are categorized as having Long COVID if they report symptoms that are new or worsened at least 90 days after an episode of COVID-19 that are not attributable to another cause; this approach is consistent with widely accepted definitions of Long COVID put forth by the World Health Organization (WHO) [17] and National Academies of Sciences, Engineering, and Medicine (NASEM) [18].

For this analysis, we studied frozen plasma samples from 15 participants in each of three clinically defined groups: fully recovered from prior SARS-CoV-2 infection (Recovered), Long COVID without neuropsychiatric symptoms (LC), and Long COVID with neuropsychiatric symptoms (NeuroLC). Neuropsychiatric symptoms included brain fog, headaches, dizziness, and difficulties with balance or vision. To compare all individuals experiencing any symptom persistence to those who recovered, we combined LC and NeuroLC participants into one group (All LC).

This research was approved by the UCSF Institutional Review Board (IRB), and all participants provided written informed consent.

### Isolation of large EVs

Plasma sample aliquots of 1 mL were precleared by centrifuging twice at 2,500 × g for 15 minutes at room temperature to pellet platelets and large cellular debris. The supernatant recovered from each spin was collected carefully to avoid disturbing the pellet. This precleared plasma was then centrifuged at 18,000 × g for 45 minutes at 4 ºC to pellet larger extracellular particles. The 18,000 × g supernatant was completely removed and stored at −80 ºC for downstream assays. The remaining pellet was resuspended in 100 µL of 0.22 µm PES-filtered PBS by rigorous pipetting after incubation at room temperature for 45 minutes and stored at −80 ºC in 25 µL aliquots for downstream nanoparticle flow cytometry analysis.

### Isolation of sEVs

Purification of sEVs was achieved by thawing a 150 µL 18,000 × g precleared plasma aliquot from each participant and loading it into a qEV 35 nm Single-Use Gen2 size-exclusion chromatography (SEC) column (IZON, #ICS35-1381) containing a Sepharose bead bed. Equilibration of the column, flushing, and purified sEV fraction collection were performed using 0.22 µm PES-filtered PBS following standard indications on the Automatic Fraction Collector (IZON AFC). A purified sEV solution of ∼370 µL was collected from each participant’s plasma sample and then stored at −80 ºC for downstream assays.

### EV quantification and characterization

Extracellular vesicles were stained with MemGlow638nm (Cytoskeleton, #MG04-10), a phospholipid bilayer dye, incubated for one hour at room temperature, and quantified using the single-particle detection Flow NanoAnalyzer U30 instrument (NanoFCM, Inc., Xiamen, China). The instrument was calibrated using standard fluorescent 250 nm silica beads of known nanoparticle concentration (NanoFCM, #QS2503). Size distribution profiles were established using the manufacturer’s calibration kits with mixtures of 150, 350, 500, and 850 nm silica beads (NanoFCM, #S17M-MV) for characterization of the 18,000 × g-enriched EVs, which we refer to as “large EVs”. These large EVs were characterized for surface mannose by staining and incubating at room temperature for one hour with the fluorescently labeled, soluble GNA (EY Labs, SKU#F-7401-2) in combination with MemGlow638nm and gated in the 100−500 nm diameter range. For flow nanoanalysis of SEC-isolated small EVs, a particle size distribution curve was established using a silica bead mixture containing 65, 100, 150, and 200 nm size nanoparticles (NanoFCM, #S16M-Exo). We refer to our SEC-isolated EVs as “small EVs” or “sEVs,” which are measured in the 40−200 nm particle diameter window and stained for the presence of phospholipid bilayer with the MemGlow638 dye.

### sEV capture assays

Two paired 150 µL aliquots of isolated sEVs from the LIINC plasma samples were compared to assess sEV capture by the GNA affinity resin. One aliquot was incubated with 20 ± 0.2 mg of GNA affinity resin (AEMD, Lot#AMI24002-023) for 1 hour at room temperature on the RotoFlex at 40 rpm, together with its paired control aliquot lacking GNA resin. After treatment, the samples were briefly spun at 1000 × g for 5 minutes to clarify the supernatant, then the sEV quantities were measured using the Flow NanoAnalyzer instrument. Differences in sEV content between the paired samples were attributed to sEV binding and capture on the GNA affinity resin.

### miRNA capture assay

330 µL aliquots of 18,000 × g-clarified plasma from All LC participants were treated with 44 ± 0.4 mg of GNA affinity resin (AEMD, Lot#AMI24002-023) by mixing for one hour at room temperature on the RotoFlex at 40 rpm together with its untreated paired control. After treatment, the samples were briefly spun for 5 minutes at 1000 × g to pellet any resin fines. RNA was isolated from 200 µL of supernatant using the miRNeasy Serum/Plasma Advanced kit (Qiagen, #217204) with the addition of 5 µL of a 200 pM mixture of three exogenous miRNAs (ath-miR159a, cel-miR-248, osa-miR414) to facilitate downstream analysis. Purified miRNA was eluted into 14 µL of RNase-free water and sent for external analysis to Bruker Spatial Biology for evaluation with the off-the-shelf NanoString nCounter miRNA expression panel (Human v3: CSO-MIR3-12), which measures the content of 827 different human miRNAs as well as the exogenous miRNA controls. Data were collected with the nCounter MAX Analysis system.

### NanoString data analysis

The NanoString nCounter analysis produced raw count data for each miRNA detected by the expression panel. The data were assessed using the nSolver 4.0 software to set a background threshold equal to the maximum output of the negative controls and normalized to the geometric mean of the three positive ligation controls and two of the exogenous spike-in miRNAs (cel-miR-248 and osa-miR414). We noted that the exogenous ath-miR159a spike-in miRNA failed to reach the detection threshold in all samples and was therefore excluded as normalization control. Additional data quality assessments identified specific samples in which either the positive ligation controls or the functional exogenous spike-in miRNAs had deficient detection signals, and these were excluded from further comparative analysis. In total, 20 paired plasma samples from All LC participants (n = 9 LC + n = 11 NeuroLC) satisfied all quality control criteria and were used to evaluate changes in miRNA content following GNA affinity resin treatment.

### Pathway activity inference

To estimate pathway activity changes downstream of miRNA depletion, we implemented a custom workflow integrating validated miRNA-mRNA interactions with footprint-based pathway models. miRNA-target interactions were obtained from miRTarBase (Release 10), using entries annotated as strong-evidence interactions [19]. For each target gene, “pseudo-expression” values were simulated under the assumption that downregulated miRNAs lead to relative de-repression of their target transcripts. When multiple miRNAs targeted the same gene, effects were aggregated using a weighted mean across interactions. These pseudo-expression profiles were analyzed using the PROGENy R package (v1.26.0, Bioconductor) to infer changes across fourteen canonical signaling pathways [20]. Pathway activity scores were obtained using all footprint genes available to maximize coverage. Directed miRNA-mRNA interactions were visualized using a layered Sugiyama layout implemented with the ggraph package. A weighted-expression heatmap was generated using the pheatmap package (v1.0.13) and a barplot of inferred pathway activity was produced with ggplot2 package (v3.5.2). All analyses were conducted in R (v4.4.1).

### Statistical analysis

Descriptive statistics were used to summarize clinical characteristics. Parametric analyses were applied to evaluate plasma EV characteristics and contents. A one-tailed t-test assuming unequal variance was used to compare EV quantities and surface glycan characteristics between the symptom groups and the control group, with p < 0.05 considered significant. Pearson’s correlation coefficient was used to examine pairwise relationships and to compare both EV quantities and mannose-positive GNA binding characteristics among distinct EV subpopulations. These analyses and graphs were generated using Excel (version 2507) and GraphPad Prism 10.6.0. To identify miRNAs significantly reduced by GNA resin treatment, normalized NanoString data were analyzed with a one-tailed t-test in R (v4.4.1) with p < 0.05 considered significant. A more conservative Hochberg correction was subsequently applied to adjust for multiple hypothesis testing and increase confidence in identifying the plasma miRNAs significantly reduced by treatment using the online calculator (https://www.multipletesting.com/analysis/) [21].

## Results

### Cohort description

Clinical characteristics were similar across the 3 groups apart from higher BMI and greater number of symptoms in the LC groups (Table 1).

**Table 1.** Demographic and clinical characteristics of participant groups. Participants fully recovered from COVID (Recovered), participants with Long COVID (LC), and symptomatic Long COVID with neurological complications (NeuroLC). All LC includes all Long COVID symptoms (LC + NeuroLC).

| Characteristic | Participant symptom group |  |  |  |
| --- | --- | --- | --- | --- |
|  | Recovered<br>(n=15) | LC<br>(n=15) | NeuroLC<br>(n=15) | All LC<br>(n=30) |
| Age, median (IQR) | 37 [30–48] | 48 [32–51] | 38 [34–46] | 40 [34–49] |
| Sex at birth |  |  |  |  |
| Female, *n* (%) | 8 (53.3) | 9 (60) | 9 (60) | 18 (60) |
| Male, *n* (%) | 7 (46.7) | 6 (40) | 6 (40) | 12 (40) |
| Race/ethnicity |  |  |  |  |
| Asian, *n* (%) | 4 (26.7) | 1 (6.7) | 1 (6.7) | 2 (6.7) |
| Hispanic/Latino, *n* (%) | 5 (33.3) | 4 (26.7) | 7 (46.7) | 11 (36.7) |
| Black/African American, *n* (%) | 0 (0) | 3 (20) | 0 (0) | 3 (10) |
| White, *n* (%) | 5 (33.3) | 7 (46.7) | 7 (46.7) | 14 (46.7) |
| Declined to answer, *n* (%) | 1 (6.7) | 0 (0) | 0 (0) | 0 (0) |
| Hospitalized during acute COVID-19 | 5 (33.3) | 4 (26.7) | 7 (46.7) | 11 (36.7) |
| BMI |  |  |  |  |
| < 25 | 8 (53.3) | 4 (26.7) | 5 (33.3) | 9 (30) |
| 25–30 | 5 (33.3) | 7 (46.7) | 6 (40) | 13 (43.3) |
| > 30 | 2 (13.3) | 4 (26.7) | 4 (26.7) | 8 (26.7) |
| Medical history |  |  |  |  |
| Autoimmune disease, *n* (%) | 1 (6.7) | 1 (6.7) | 2 (13.3) | 3 (10) |
| Cancer treated within past 2 years, *n* (%) | 0 (0) | 1 (6.7) | 0 (0) | 1 (3.3) |
| Diabetes, *n* (%) | 3 (20) | 1 (6.7) | 1 (6.7) | 2 (6.7) |
| Heart attack or heart failure, *n* (%) | 0 (0) | 0 (0) | 0 (0) | 0 (0) |
| Hypertension or high blood pressure, *n* (%) | 0 (0) | 3 (20) | 3 (20) | 6 (20) |
| Lung disease, *n* (%) | 2 (13.3) | 5 (33.3) | 1 (6.7) | 6 (20) |
| Kidney disease, *n* (%) | 0 (0) | 1 (6.7) | 0 (0) | 1 (3.3) |
| Long COVID symptoms, median (IQR) | 0 [0–0] | 3 [1–3] | 5 [3–8] | 3 [3–5] |
| Days post symptom onset, median (IQR) | 72 [68–88] | 77 [62–86] | 78 [73–81] | 78 [66–83] |

Abbreviations: BMI, body mass index (weight in kg divided by the height in meters squared); IQR, interquartile range (25<sup>th</sup>–75<sup>th</sup> percentile); *n* (%), number and percentage of participants in the corresponding group.

### Plasma EV characterization

Plasma large EVs (100–500 nm) displayed substantial variability in individual sample concentration (Figure 1A). On average, the NeuroLC plasma had 64% higher concentrations of these vesicles than LC and Recovered plasma; however, this difference was not statistically significant (Figure 1B). Small EVs (40–200 nm) isolated from plasma also displayed considerable individual sample variability (Figure 1C), with average quantities highest in the NeuroLC group, although this increase was not statistically significant (Figure 1D).

**Figure 1.**
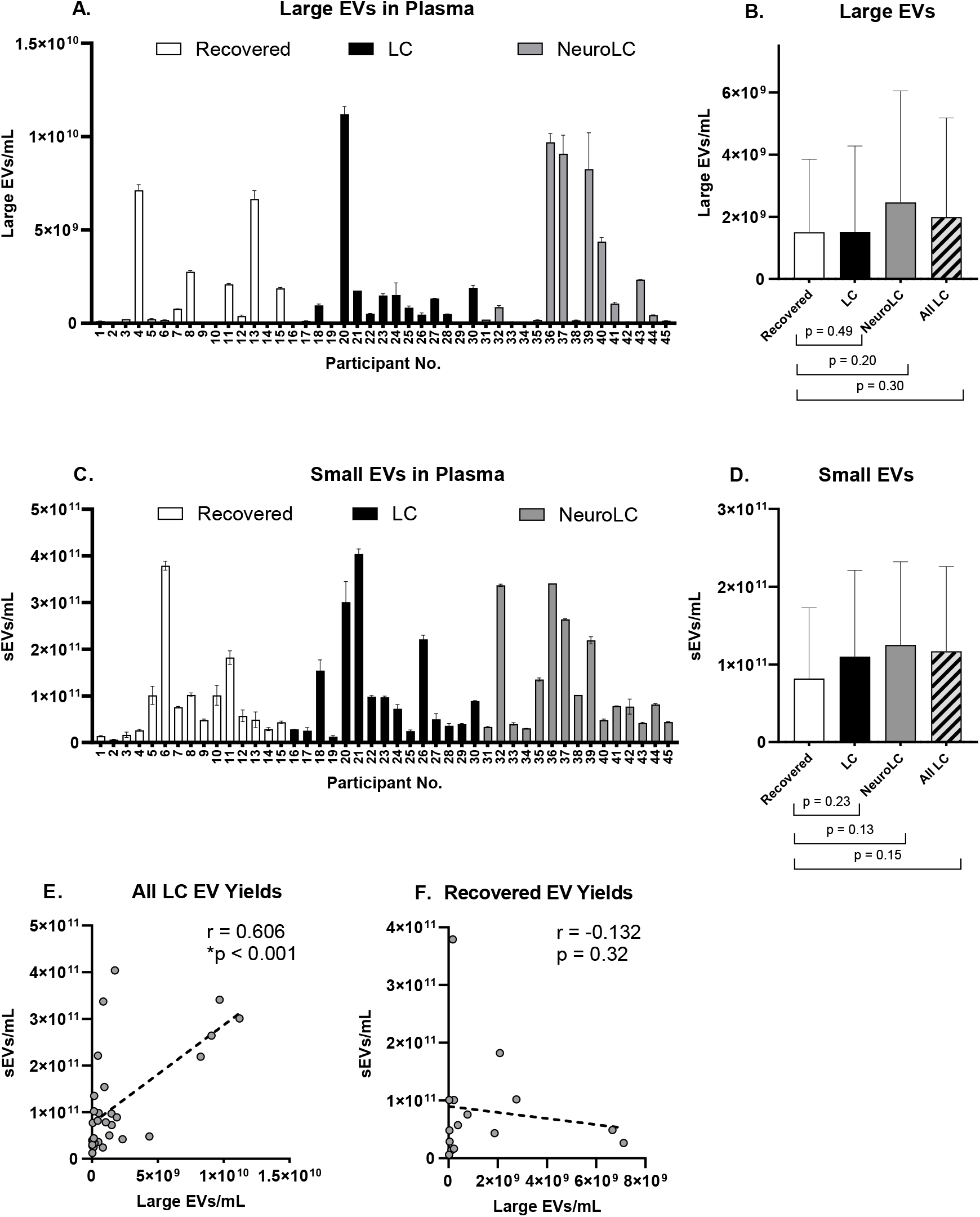
Plasma extracellular vesicle yields in the post-acute phase of COVID-19. A) Large EV yields (100−500 nm). Data shown represents the average and SD of two technical replicates; Average ± SD quantities of large EVs per symptom group (n = 15); C) small EV yields (40−200 nm). Data shows the average and SD of two technical replicates; D) Average ± SD quantities of sEVs measured in each symptom group (n = 15); E) Correlation of large and small EV quantities isolated from All LC (r = 0.606, *p < 0.001, n = 30); F) and from Recovered plasmas (r = –0.132, p = 0.32, n = 15).

Evidence of increased circulating EV production in Long COVID plasma was observed across multiple EV subpopulation size ranges. When we examined the association between quantities of large and small EVs in All LC we detected a significant positive correlation between concentrations of these two EV subpopulations (Figure 1E). In contrast, no such association was observed among EVs from fully recovered participants as no relationship was detected among concentrations of large and small EVs isolated from Recovered plasma (Figure 1F). These findings suggest there may be a disease-associated EV production mechanism affecting multiple subpopulations.

In order to determine if there were differences in the glycosylation patterns of EVs in the distinct Long COVID symptom groups, we evaluated concentrations of mannose-positive large EVs (100−500 nm) in these plasma samples. We observed a large range of individual variability in the quantities of mannose-positive large EVs (Figure 2A), with average quantities two-fold significantly higher in All LC groups (Figure 2B). Although binding of fluorescently labeled GNA to mannose glycans on the surface of smaller EVs (<100 nm) did not produce a signal strong enough to quantify, capture of sEVs to the GNA affinity resin could be tracked. On average, we observed 31-42% increased sEV capture by GNA affinity resin treatment of All LC groups compared with the control (Recovered), although these differences were not significant (Figure 2C). Importantly, a significant positive correlation was observed between mannose-positive large EV quantities and sEV capture by GNA resin treatment in All LC samples (Figure 2D). In contrast, no association was found between mannose-positive large EV abundance and GNA resin capture of sEVs isolated from plasma of Recovered controls (Figure 1F).

**Figure 2.**
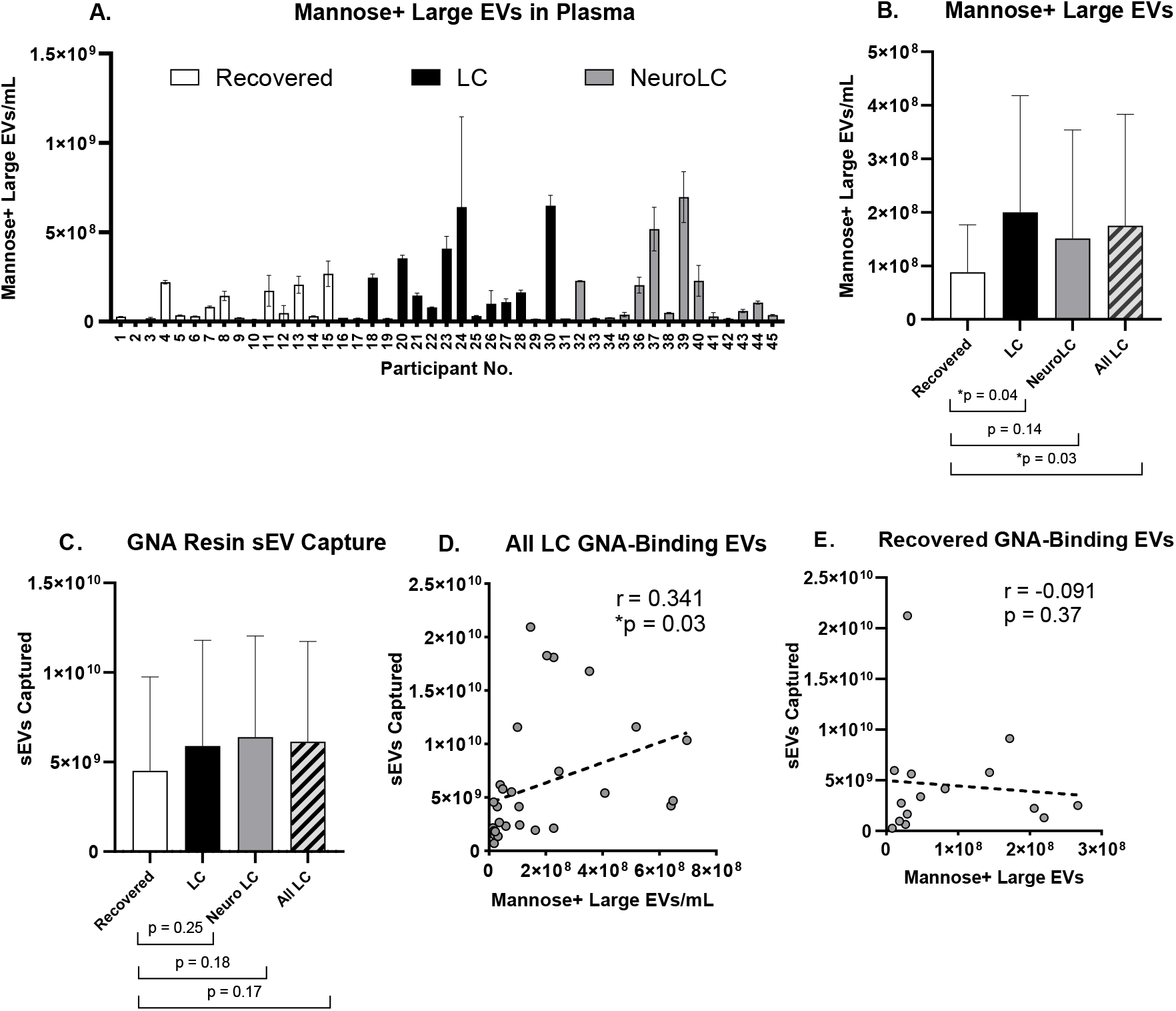
Increased mannose glycosylation in symptomatic plasma EVs improves GNA-based capture. A) Abundance of mannose-positive large EVs (100−500 nm). Data shown represents the average and SD of two technical replicates; B) Average quantities of mannose-positive large EVs measured in each symptom group (n = 15, except for All LC where n = 30). Comparisons to control (Recovered) group show statistical differences (*p < 0.05). C) Average quantities of sEVs (40−200 nm) removed by GNA affinity resin treatment by symptom group (n = 15, except for All LC where n = 30), D) Correlation between sEV capture on GNA affinity resin and mannose-positive EV abundance in All LC (r = 0.341, *p < 0.05, n = 30); E) and in Recovered samples (r = –0.091, p = 0.37, n = 15).

### Analysis of EV-associated miRNA cargo

To characterize miRNA cargo of sEVs removed by GNA affinity resin treatment, we analyzed changes to miRNA content of All LC plasma samples (n = 20) after incubation with GNA affinity resin using NanoString nCounter technology. Of the 827 human miRNAs of biological relevance that can be evaluated using the off-the-shelf NanoString miRNA panel, 358 were detected in quantities above the background threshold. Approximately 36% of these miRNAs (128/358) were found to be significantly removed by GNA affinity resin when pre-vs. post-treatment levels were compared using a one-tailed t-test (p < 0.05; Supplementary Table 1). To eliminate large-scale false discovery, a multiple hypothesis testing correction was applied using the Hochberg adjustment method available through this online analytics tool https://multipletesting.com [21]. After performing the Hochberg multiple hypothesis testing correction, seven miRNAs (hsa-miR-374a-5p, hsa-miR-640, hsa-miR-301b-3p, hsa-miR-1272, hsa-miR-3613-3p, hsa-miR-4531, hsa-miR-874-3p) were identified as being significantly reduced by GNA affinity resin treatment (Table 2).

**Table 2.** miRNAs significantly depleted by GNA affinity resin treatment in All LC plasma samples.

| miRNA | Accession # | Log2 Fold Change | Percent Reduction |
| --- | --- | --- | --- |
| hsa-miR-374a-5p | MIMAT0000727 | -0.26 | -19.9% |
| hsa-miR-640 | MIMAT0003310 | -0.27 | -20.3% |
| hsa-miR-301b-3p | MIMAT0004958 | -0.29 | -21.9% |
| hsa-miR-1272 | MIMAT0005925 | -0.29 | -22.0% |
| hsa-miR-3613-3p | MIMAT0017991 | -0.38 | -30.0% |
| hsa-miR-4531 | MIMAT0019070 | -0.26 | -19.4% |
| hsa-miR-874-3p | MIMAT0004911 | -0.23 | -17.3% |

To explore the potential effects of depletion of these 7 miRNAs, we selected a methodology to expand our understanding of their mRNA targets. We used miRTarBase to retrieve evidence of experimentally validated miRNA-target interactions. As shown in Figure 3A, we identified a total of 32 gene transcripts targeted by one or more of these miRNAs. We then used the PROGENy pathway analysis tool to gain insight into how 14 well-established canonical signaling pathways are affected by perturbations in activity of these genes. For this analysis, we assumed that the reduction of a miRNA suppressor could result in an equal proportional uptick in target gene expression. A PROGENy weighted-expression heatmap was created to depict how these potential escalations in expression of the 32 gene targets would affect the core signaling pathways (Figure 3B). Lastly, an activity score assigned to each pathway demonstrates that the signaling pathway inferred to have the most enhanced upregulation as a result of GNA resin treatment is the Estrogen pathway, while the most downregulated is the JAK-STAT (Figure 3C).

**Figure 3.**
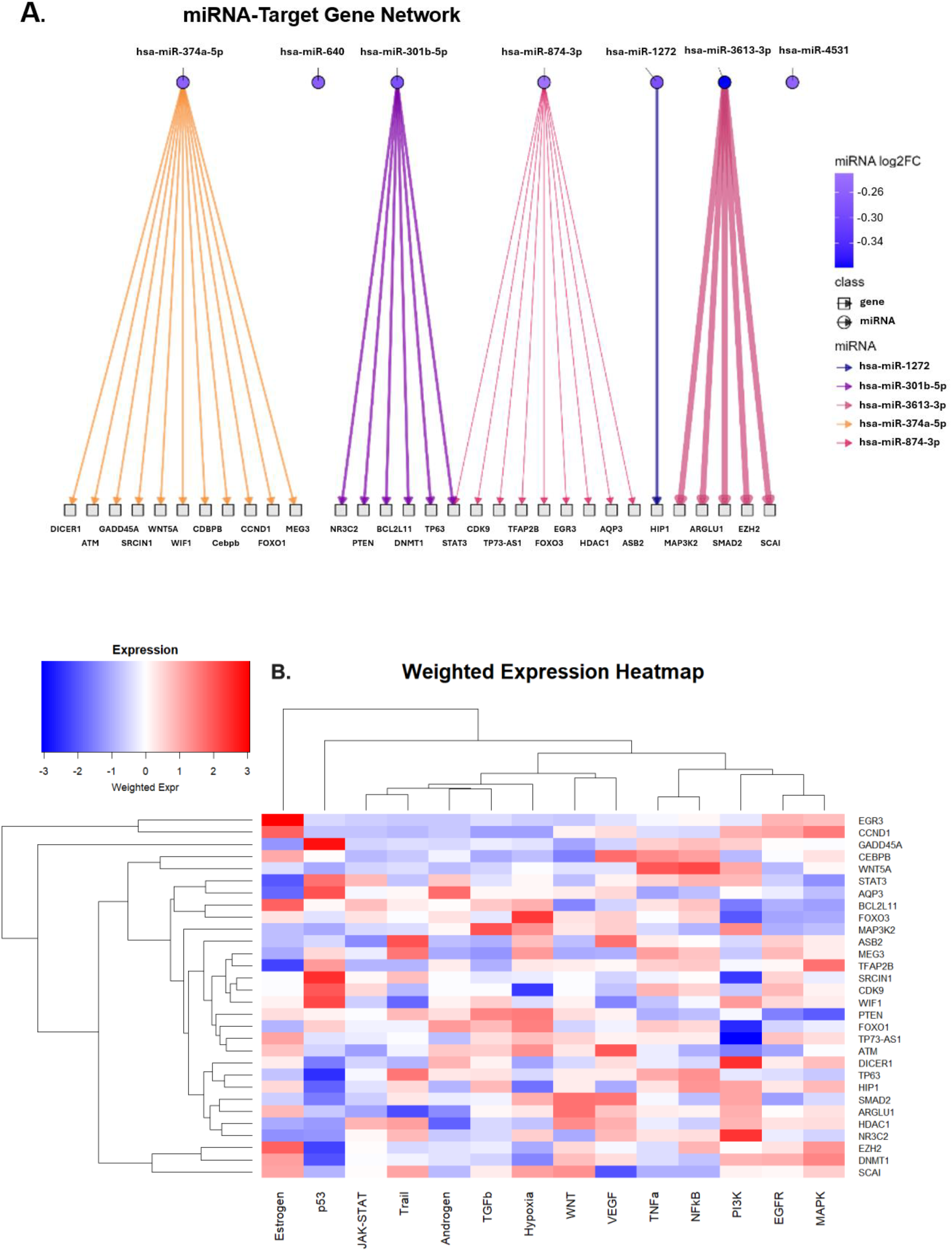
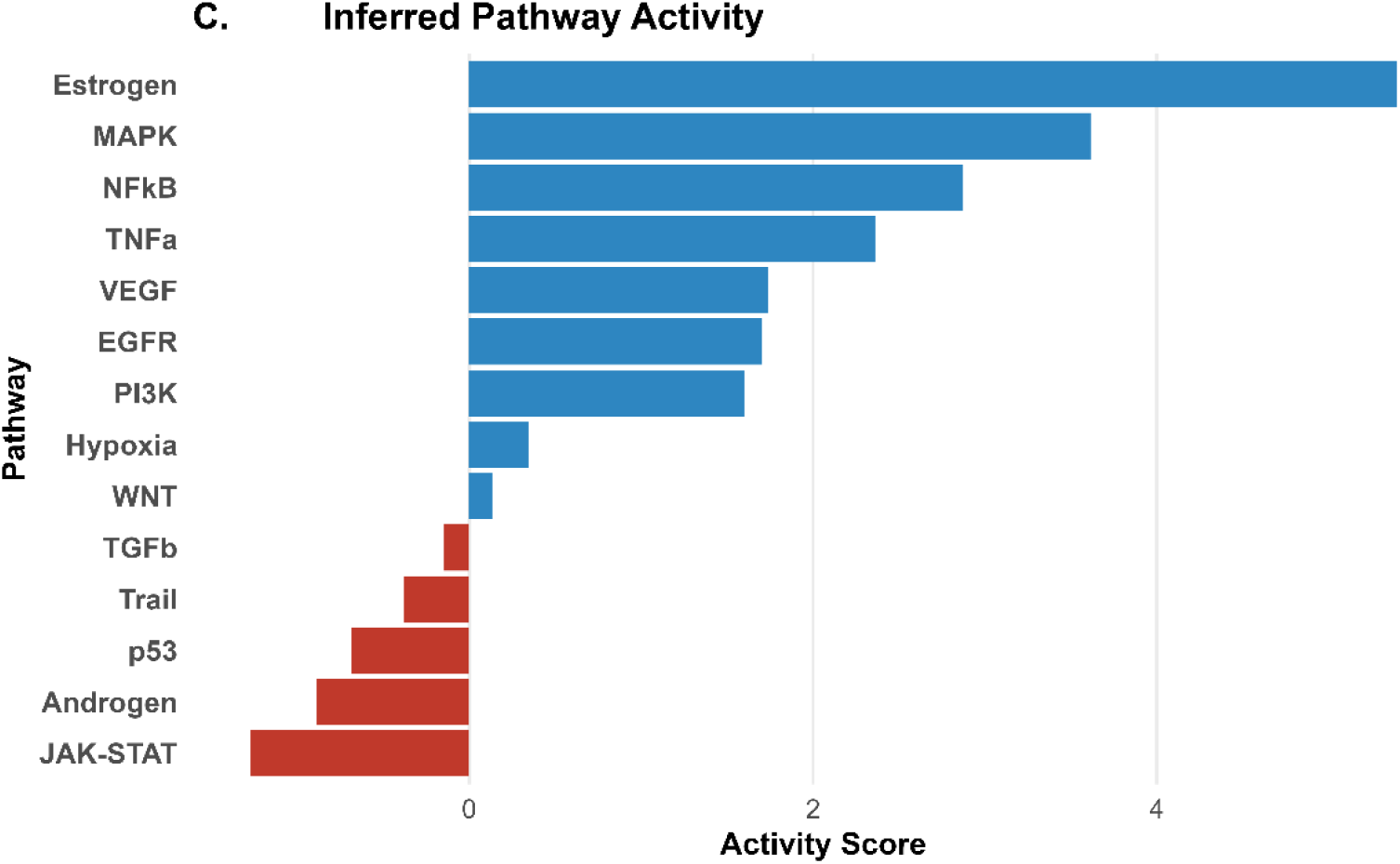
Inferential miRNA pathway analysis. A) Sugiyama Framework plot depicting experimentally validated target genes of the 7 miRNAs significantly reduced by GNA resin treatment. Line thickness corresponds to the magnitude of the miRNA log2(fold change). B) Weighted Expression Heatmap depicting relevance of potential target gene changes on signaling pathways (blue = suppressed, white = unchanged, red = de-repressed). C) Inferred Activity Score of canonical signaling pathway modulations.

## Discussion

We found substantial variability in plasma EV populations during the post-acute phase of COVID-19, with some EV characteristics more common among individuals with Long COVID than those who fully recovered from their infection. This suggests that disease-associated EV production mechanisms may be relevant in this patient population. Compared to those who fully recovered, large EVs in those with Long COVID had distinct surface glycosylation characteristics with significantly higher mannose content. Finally, we observed that these EVs can be captured using GNA, a mannose-specific binding lectin, suggesting a potential therapeutic approach that warrants further study.

Even though there is no existing literature assessing mannosylation on EVs in acute SARS-CoV-2 infection or Long COVID, there is evidence suggesting that the glycosylation pathway of the host during acute infection is altered. SARS-CoV-2 spike contains many glycosylation sites, several of which consistently retain high-mannose glycans [22]. Viral entry into ACE2-expressing host cells requires spike, and both viral and host glycoproteins utilize the endoplasmic reticulum (ER) and Golgi glycosylation machinery [23, 24]. Genome-wide CRISPR screens and perturbation studies have identified N-glycosylation enzymes as essential host dependency factors for SARS-CoV-2 infection [25, 26]. Clinical glycomics analyses of acute COVID-19 further demonstrate increased high-mannose content on immunoglobulins, particularly IgM, which correlates with disease severity [27]. This offers evidence that the glycosylation pathway plays an important role in acute infection and is also likely to be altered. So far, this alteration remains to be proven in Long COVID. Our findings of increased mannosylation on plasma EVs is suggestive of ongoing glycosylation pathway abnormalities that may play a role in Long COVID pathology.

In Long COVID, circulating dysregulated miRNA profiles can also contribute to pathogenic processes including inflammation and immune dysregulation [28]. Similarly, in acute COVID-19, specific miRNA enriched within circulating extracellular vesicles (EV-miRNA) have been shown to participate in proinflammatory and prothrombotic complications [29]. In this study, we wanted to understand the nature of the plasma EV-miRNA cargo within the mannose-positive EVs that can be selectively targeted and eliminated by capture on the GNA affinity resin. As such, we compared the circulating miRNA profile in Long COVID plasma before and after incubation on the GNA affinity resin, to identify miRNA significantly reduced by this treatment. Our miRNA analysis provides insight into the potential clinical benefits of mannose-positive EV capture from Long COVID patient plasma.

The seven miRNAs significantly reduced by GNA allowed us to infer the post-treatment changes in activity of 14 fundamental canonical pathways, which suggest a favorable rebalancing of immune and tissue-repair networks (Table 2, Figure 3). The JAK-STAT pathway alteration, inferred to be de-activated with greatest magnitude, may alleviate sustained symptoms caused by ongoing interferon signaling [30]. Inhibition of this pathway is now of great interest, and our findings provide additional rationale for trials of small molecule JAK inhibitors for patients experiencing Long COVID–related neurocognitive and cardiopulmonary symptoms that are now underway (NCT05858515, NCT06597396, NCT06928272). Furthermore, the observed activation of Estrogen, VEGF, EGFR, and P13K pathways may remediate the reported endothelial dysfunction in Long COVID by enhancing microvascular support, nitric oxide-mediated vasodilation, and endothelial repair [31-33]. On the other hand, activation of pathways such as NF-kB, MAPK, and TNF-alpha is generally linked to the amplification of inflammatory cytokines [34]. Therefore, GNA affinity resin inferred activation of such pathways might seem counterintuitive given that subsets of Long COVID patients have persistent inflammatory activity. However, multi-omics data demonstrates that Long COVID is quite heterogeneous, with patient subsets having suppressed NF-kB signaling and impaired immune responsiveness [35]. For these patients, an uptick of the NF-kB, MAPK, and TNF-alpha pathways could have a restorative effect of the innate signaling circuits that would otherwise be dampened.

Beyond small molecule therapeutics, the observation that sEV capture was significantly associated with mannose-positive large EV measurements in All LC participants suggests that there is a mannose glycome alteration affecting multiple EV subpopulations, making a GNA lectin-based treatment a highly promising approach for EV capture from Long COVID patient plasma. Removal of these EVs might have important implications given the recent findings suggesting that disease relevant EVs may be clinically meaningful, since EVs have been implicated as carriers of viral antigens and pro-inflammatory signals in Long COVID [10, 36]. Targeted removal of these vesicles could therefore represent a promising adjunctive strategy to reduce persistent immune activation and symptom burden.

This study has several limitations. The small sample size in this exploratory pilot study limits the statistical power of analysis potentially masking additional discoveries. The study was cross-sectional, and the timing of samples studied was relatively early in the post-acute phase, slightly before the recently defined 90-day threshold for Long COVID. The NanoString nCounter platform utilizes an 827 predetermined human miRNA panel, excluding potential treatment relevant miRNA changes not part of the repertoire. The removal effect of those miRNAs on signaling pathways was computationally inferred using PROGENy. In this process, we mapped the miRNA-mRNA interactions from miRTarBase using only strong-evidence interactions, thus increasing the confidence of our inferred miRNA effects. Nevertheless, this approach might also be excluding less well-characterized miRNA-mRNA interactions. We also did not interrogate the mannosylated EVs for SARS-CoV-2 particles, autoantibodies or reactivated EBV which have been implicated in the pathogenesis of Long COVID. These experiments would be the next pre-clinical step in considering the potential utility of the Hemopurifier in Long COVID. Ultimately, a clinical trial interrogating the ability of the Hemopurifier® to resolve the pathology and symptomology of Long COVID patients is necessary to confirm these inferred pathway changes.

## Acknowledgements

We are grateful to the LIINC study participants and to the clinical staff who provided care to these individuals during their acute illness and during their recovery. We also acknowledge the current and former LIINC clinical study team members who collected the clinical data and biospecimens. M.J.P. is supported on K23AI157875. The LIINC clinical core is supported by the PolyBio Research Foundation, with additional funding from R01AI141003 (to T.J.H.) and R01NS136197 (to M.J.P.).

## Ethics

The LIINC program was approved by the UCSF Institutional Review Board. All participants provided written informed consent.

## Author Contributions

Conceptualization, S.P.L. and M.J.P.; methodology, M.P. and R.d.N.C.; validation, M.P. and R.d.N.C.; formal analysis, M.P. and R.d.N.C.; investigation, M.P. and R.d.N.C.; experimental data curation, M.P.; clinical data curation, T.D., P.C., and E.A.F.; LIINC cohort study design, J.D.K., J.N.M., S.G.D., T.J.H., and M.J.P.; resources, S.P.L. and M.J.P.; visualization, M.P.; supervision, R.d.N.C. and S.P.L.; project administration, S.P.L.; writing—original draft preparation, M.P. and R.d.N.C.; writing—review and editing, all authors. All authors have read and agreed to the published version of the manuscript.

## Disclosures

This study was funded by Aethlon Medical, Inc., manufacturer of the Hemopurifier. Authors R.d.N.C, M.P. and S.P.L. are employees of Aethlon Medical, Inc.

The authors used ChatGPT to assist in generating R code and drafting relevant descriptive methods, with all content subsequently reviewed and edited by the authors for accuracy and clarity.

