## Supplemental Table 1 for "Increased mannosylation of extracellular vesicles in Long COVID plasma provides a potential therapeutic target for *Galanthus nivalis* agglutinin (GNA) affinity resin"

**Supplementary Table 1. List of miRNAs reduced by GNA resin treatment in All LC plasma samples.**

| miRNA | Accession # | $Log_2$ (fold change) | p value |
|---|---|---|---|
| hsa-miR-374a-5p | MIMAT0000727 | -0.26 | 0.002 |
| hsa-miR-640 | MIMAT0003310 | -0.27 | 0.002 |
| hsa-miR-301b-3p | MIMAT0004958 | -0.29 | 0.003 |
| hsa-miR-1272 | MIMAT0005925 | -0.29 | 0.005 |
| hsa-miR-3613-3p | MIMAT0017991 | -0.38 | 0.005 |
| hsa-miR-4531 | MIMAT0019070 | -0.26 | 0.005 |
| hsa-miR-874-3p | MIMAT0004911 | -0.23 | 0.007 |
| hsa-miR-320e | MIMAT0015072 | -0.27 | 0.007 |
| hsa-miR-590-5p | MIMAT0003258 | -0.23 | 0.007 |
| hsa-miR-455-3p | MIMAT0004784 | -0.23 | 0.007 |
| hsa-miR-371a-5p | MIMAT0004687 | -0.41 | 0.007 |
| hsa-miR-1183 | MIMAT0005828 | -0.23 | 0.007 |
| hsa-miR-133a-3p | MIMAT0000427 | -0.31 | 0.008 |
| hsa-miR-210-3p | MIMAT0000267 | -0.38 | 0.008 |
| hsa-miR-652-5p | MIMAT0022709 | -0.23 | 0.008 |
| hsa-miR-2110 | MIMAT0010133 | -0.30 | 0.008 |
| hsa-miR-4707-5p | MIMAT0019807 | -0.24 | 0.008 |
| hsa-miR-502-3p | MIMAT0004775 | -0.22 | 0.008 |
| hsa-miR-574-3p | MIMAT0003239 | -0.27 | 0.009 |
| hsa-miR-1286 | MIMAT0005877 | -0.23 | 0.009 |
| hsa-miR-587 | MIMAT0003253 | -0.21 | 0.009 |
| hsa-miR-513a-3p | MIMAT0004777 | -0.27 | 0.010 |
| hsa-miR-182-3p | MIMAT0000260 | -0.22 | 0.010 |
| hsa-miR-1257 | MIMAT0005908 | -0.22 | 0.010 |
| hsa-miR-20a-5p+hsa-miR-20b-5p | MIMAT0000075 | -0.23 | 0.010 |
| hsa-miR-1304-5p | MIMAT0005892 | -0.35 | 0.010 |
| hsa-miR-3180-3p | MIMAT0015058 | -0.25 | 0.010 |
| hsa-miR-219b-3p | MIMAT0019748 | -0.24 | 0.010 |
| hsa-miR-328-5p | MIMAT0026486 | -0.28 | 0.010 |
| hsa-miR-548ak | MIMAT0019013 | -0.23 | 0.011 |
| hsa-miR-556-3p | MIMAT0004793 | -0.23 | 0.011 |
| hsa-miR-520f-3p | MIMAT0002830 | -0.21 | 0.011 |
| hsa-miR-128-1-5p | MIMAT0026477 | -0.22 | 0.011 |
| hsa-miR-135b-5p | MIMAT0000758 | -0.22 | 0.011 |
| hsa-miR-549a | MIMAT0003333 | -0.34 | 0.012 |

| | | | |
|---|---|---|---|
| hsa-miR-34a-5p | MIMAT0000255 | -0.23 | 0.012 |
| hsa-miR-1537-3p | MIMAT0007399 | -0.26 | 0.012 |
| hsa-miR-660-5p | MIMAT0003338 | -0.23 | 0.012 |
| hsa-miR-19b-3p | MIMAT0000074 | -0.29 | 0.012 |
| hsa-miR-412-3p | MIMAT0002170 | -0.25 | 0.013 |
| hsa-miR-1307-5p | MIMAT0022727 | -0.21 | 0.013 |
| hsa-miR-190a-3p | MIMAT0026482 | -0.21 | 0.013 |
| hsa-miR-503-3p | MIMAT0022925 | -0.21 | 0.013 |
| hsa-miR-6511a-3p | MIMAT0025479 | -0.21 | 0.013 |
| hsa-miR-499b-3p | MIMAT0019898 | -0.21 | 0.013 |
| hsa-miR-23c | MIMAT0018000 | -0.41 | 0.014 |
| hsa-miR-4755-5p | MIMAT0019895 | -0.22 | 0.014 |
| hsa-miR-23b-3p | MIMAT0000418 | -0.27 | 0.014 |
| hsa-miR-1973 | MIMAT0009448 | -0.25 | 0.014 |
| hsa-miR-1307-3p | MIMAT0005951 | -0.21 | 0.014 |
| hsa-miR-509-3p | MIMAT0002881 | -0.21 | 0.014 |
| hsa-miR-19a-3p | MIMAT0000073 | -0.22 | 0.014 |
| hsa-miR-3185 | MIMAT0015065 | -0.21 | 0.015 |
| hsa-miR-570-3p | MIMAT0003235 | -0.25 | 0.015 |
| hsa-miR-936 | MIMAT0004979 | -0.21 | 0.015 |
| hsa-miR-18b-5p | MIMAT0001412 | -0.21 | 0.016 |
| hsa-miR-3161 | MIMAT0015035 | -0.21 | 0.016 |
| hsa-miR-487b-3p | MIMAT0003180 | -0.21 | 0.016 |
| hsa-miR-499a-5p | MIMAT0002870 | -0.21 | 0.016 |
| hsa-miR-524-3p | MIMAT0002850 | -0.21 | 0.016 |
| hsa-miR-3613-5p | MIMAT0017990 | -0.21 | 0.016 |
| hsa-miR-631 | MIMAT0003300 | -0.21 | 0.016 |
| hsa-miR-4787-5p | MIMAT0019956 | -0.21 | 0.016 |
| hsa-miR-455-5p | MIMAT0003150 | -0.27 | 0.016 |
| hsa-miR-1203 | MIMAT0005866 | -0.24 | 0.016 |
| hsa-miR-876-5p | MIMAT0004924 | -0.22 | 0.016 |
| hsa-miR-217 | MIMAT0000274 | -0.25 | 0.017 |
| hsa-miR-548m | MIMAT0005917 | -0.34 | 0.018 |
| hsa-miR-423-3p | MIMAT0001340 | -0.21 | 0.018 |
| hsa-let-7c-5p | MIMAT0000064 | -0.21 | 0.018 |
| hsa-miR-125b-5p | MIMAT0000423 | -0.21 | 0.018 |
| hsa-miR-365b-5p | MIMAT0022833 | -0.21 | 0.018 |
| hsa-miR-221-3p | MIMAT0000278 | -0.21 | 0.018 |

| | | | |
|---|---|---|---|
| hsa-miR-302a-5p | MIMAT0000683 | -0.21 | 0.018 |
| hsa-miR-1193 | MIMAT0015049 | -0.20 | 0.019 |
| hsa-miR-1205 | MIMAT0005869 | -0.20 | 0.019 |
| hsa-miR-1258 | MIMAT0005909 | -0.20 | 0.019 |
| hsa-miR-1264 | MIMAT0005791 | -0.20 | 0.019 |
| hsa-miR-1322 | MIMAT0005953 | -0.20 | 0.019 |
| hsa-miR-143-3p | MIMAT0000435 | -0.20 | 0.019 |
| hsa-miR-224-5p | MIMAT0000281 | -0.20 | 0.019 |
| hsa-miR-302f | MIMAT0005932 | -0.20 | 0.019 |
| hsa-miR-3180 | MIMAT0018178 | -0.20 | 0.019 |
| hsa-miR-516a-3p+hsa-miR-516b-3p | MIMAT0006778 | -0.20 | 0.019 |
| hsa-miR-518f-3p | MIMAT0002842 | -0.20 | 0.019 |
| hsa-miR-541-3p | MIMAT0004920 | -0.20 | 0.019 |
| hsa-miR-542-5p | MIMAT0003340 | -0.20 | 0.019 |
| hsa-miR-564 | MIMAT0003228 | -0.20 | 0.019 |
| hsa-miR-644a | MIMAT0003314 | -0.20 | 0.019 |
| hsa-miR-940 | MIMAT0004983 | -0.20 | 0.019 |
| hsa-miR-1224-3p | MIMAT0005459 | -0.20 | 0.019 |
| hsa-miR-1261 | MIMAT0005913 | -0.20 | 0.019 |
| hsa-miR-141-3p | MIMAT0000432 | -0.20 | 0.019 |
| hsa-miR-147b | MIMAT0004928 | -0.20 | 0.019 |
| hsa-miR-506-3p | MIMAT0002878 | -0.20 | 0.019 |
| hsa-miR-612 | MIMAT0003280 | -0.28 | 0.019 |
| hsa-miR-2113 | MIMAT0009206 | -0.21 | 0.019 |
| hsa-miR-181a-5p | MIMAT0000256 | -0.21 | 0.020 |
| hsa-miR-449a | MIMAT0001541 | -0.20 | 0.020 |
| hsa-miR-411-5p | MIMAT0003329 | -0.23 | 0.020 |
| hsa-miR-1245a | MIMAT0005897 | -0.25 | 0.021 |
| hsa-miR-502-5p | MIMAT0002873 | -0.32 | 0.022 |
| hsa-miR-517c-3p+hsa-miR-519a-3p | MIMAT0002866 | -0.29 | 0.023 |
| hsa-miR-556-5p | MIMAT0003220 | -0.22 | 0.023 |
| hsa-miR-376a-2-5p | MIMAT0022928 | -0.19 | 0.024 |
| hsa-miR-128-3p | MIMAT0000424 | -0.19 | 0.025 |
| hsa-miR-134-3p | MIMAT0026481 | -0.24 | 0.025 |
| hsa-miR-208a-3p | MIMAT0000241 | -0.25 | 0.026 |
| hsa-miR-92b-3p | MIMAT0003218 | -0.19 | 0.026 |
| hsa-miR-6720-3p | MIMAT0025851 | -0.27 | 0.026 |
| hsa-miR-345-3p | MIMAT0022698 | -0.22 | 0.027 |

| | | | |
|---|---|---|---|
| hsa-miR-106a-5p+hsa-miR-17-5p | MIMAT0000103 | -0.24 | 0.028 |
| hsa-miR-548al | MIMAT0019024 | -0.26 | 0.030 |
| hsa-miR-603 | MIMAT0003271 | -0.65 | 0.030 |
| hsa-miR-302a-3p | MIMAT0000684 | -0.19 | 0.031 |
| hsa-miR-656-3p | MIMAT0003332 | -0.18 | 0.031 |
| hsa-miR-342-3p | MIMAT0000753 | -0.19 | 0.033 |
| hsa-miR-24-3p | MIMAT0000080 | -0.24 | 0.035 |
| hsa-miR-4461 | MIMAT0018983 | -0.27 | 0.037 |
| hsa-miR-582-5p | MIMAT0003247 | -0.25 | 0.039 |
| hsa-miR-29c-3p | MIMAT0000681 | -0.18 | 0.043 |
| hsa-miR-29a-3p | MIMAT0000086 | -0.22 | 0.043 |
| hsa-miR-299-5p | MIMAT0002890 | -0.25 | 0.045 |
| hsa-miR-548aa+hsa-miR-548t-3p | MIMAT0018447 | -0.39 | 0.046 |
| hsa-miR-496 | MIMAT0002818 | -0.28 | 0.047 |
| hsa-miR-1278 | MIMAT0005936 | -0.20 | 0.048 |
| hsa-miR-548v | MIMAT0015020 | -0.29 | 0.048 |
